# PanACRpred: Predicting Accessible Chromatin Regions in Pangenomes using Motif Chaining

**DOI:** 10.64898/2026.02.05.703812

**Authors:** Madelyn Warr, Trung Dinh, Bella Root, Elyse Onstott, Kevin Yu, Joann Mudge, Thiruvarangan Ramaraj, Indika Kahanda, Brendan Mumey

## Abstract

In this work, we investigate using motif subsequence features to predict whether a genomic region is accessible to regulatory proteins, i.e. an accessible chromatin region (ACR), enabling transcription of associated genes. We focus on plants, whose agricultural and ecological importance make them interesting and important organisms to study, and whose complex genomes provide important stress tests for our algorithm. We show that motif sequence similarity as found by co-linear chaining can be used in combination with machine learning models to effectively predict ACRs in genome assemblies.

## 1. Introduction

As DNA sequencing technologies and assembly algorithms have improved, many high-quality genome assemblies are often being generated in a single species [9,20]. This has allowed us to move away from analyzing the genetic variation in a single species through comparison of sparse sequence data from multiple individuals back to a single reference, which introduces bias. Instead, we can now create pangenome data structures that uniformly capture genetic variation from individuals to enable capture and analysis of all the genetic variation in a species. While the availability of multiple genomes per species is relatively common, other sequencing data types are lagging behind.

In addition to understanding the genetic variation present in a species, it is also important to understand how that variation is expressed. *ATAC-seq* is a technology that sequences regions of the genome that are accessible to proteins, known as accessible chromatin regions (ACRs). These regions bind regulatory factors, thus marking clusters of *cis-regulatory elements* (CREs) such as promoters and enhancers that are actively driving transcription of their corresponding genes [14,31,33]. CREs are noncoding DNA regions in, around, or distal to genes that affect expression timing, tissue-specificity, and environmental response [18,3,4,21]. CREs affect expression by binding transcription factors, small RNAs, enzymes affecting DNA modifications, and/or modified histones [28], bringing them into physical contact with the gene they control, often through chromosome looping [32,26].

ATAC-seq is a powerful tool for investigating the role of ACRs and CREs in regulating gene expression. Additionally, ATAC-seq can be used to identify the short conserved DNA sequences that characterize CREs referred to as *motifs*. However, ATAC-seq data are still limited, often focused on multiple tissues of the reference genome, rather than being assessed across multiple individuals. Therefore, in a pangenomic setting, where genome assemblies of many individuals are available, ATAC-seq data and the accessible chromatin regions they identify may only be available for one or two lines.

In this work, we investigate whether the CRE content in a region can be used to predict chromatin accessibility in a given condition. We focus on plants, whose agricultural and ecological importance, combined with their complex genomes make them interesting and important organisms to study [27]. We show that limited ATAC-seq data combined with CRE information can be used to predict ACRs in genome assemblies. Accessible chromatin regions associated with the same genes have some short regions of CRE conservation [16,15,30]. As a first step toward chaining CREs across pangenomes (Fig. 1), we quantify CRE similarity between genome regions using *co-linear chaining* [17,8] to identify candidate chromatin-accessible regions by identifying motifs shared between a known accessible region and related regions. Finally, the chaining scores against known ACRs are used as a *feature vector* to predict accessibility of unknown regions. We show this strategy is effective at accurately predicting novel ACR regions.

**Fig. 1.**
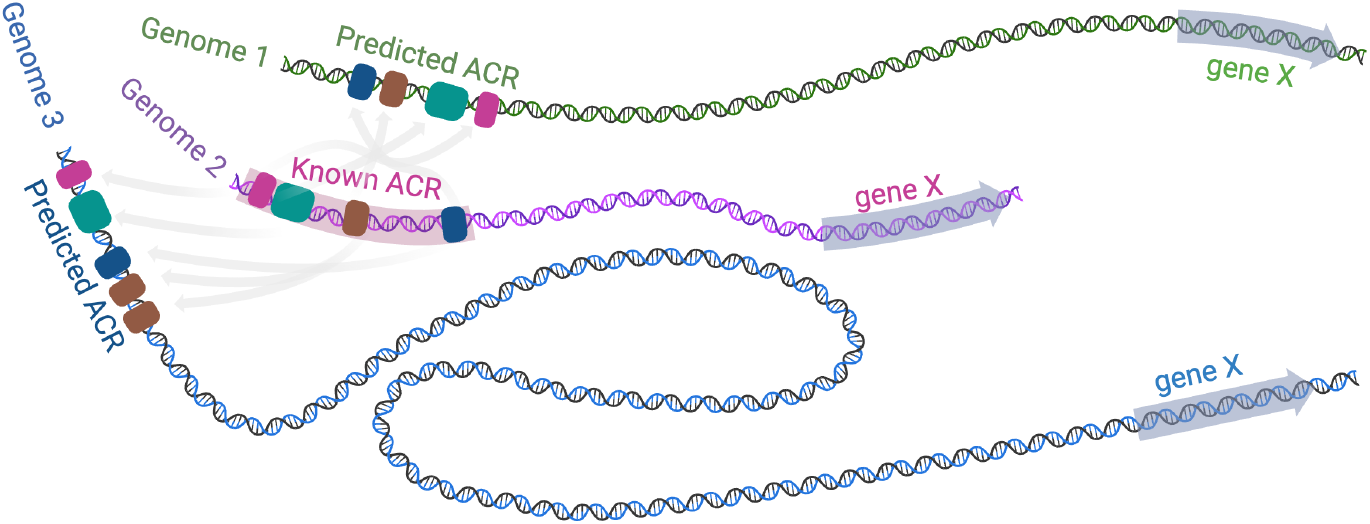
Co-linear chaining can quickly find motif subsequence matching scores between known ACRs and candidate regions; these scores are used to train a machine learning classifier to predict accessibility of candidate regions in a pangenomic data set. Created by BioRender.

## 2 Related Work

Early machine learning methods to predict regulatory elements such as gkm-SVM [5] relied on hand-crafted features, primarily k-mers. DeepSEA [34] was one of the first deep learning models, which used a Convolutional Neural Network (CNN) to predict chromatin effects directly from raw DNA sequences, outperforming SVMs by capturing hierarchical features. Then, researchers began using specialized architectures to capture the “grammar” of the genome, which introduced (a) Basset [11], which applies CNNs specifically to ATAC-seq data across multiple cell types simultaneously, (b) Deopen [13], which combined CNNs with k-mer features to improve accuracy in predicting continuous DNase-seq signals, and (c) DanQ [24], which combined CNNs with LSTMs (Long Short-Term Memory) to capture both local motifs and their long-range spatial relationships. Current State-of-the-Art models focus on high-resolution (base-pair level) and long-range context. For example, Basenji [10] uses dilated convolutions allowing the model to look at sequences up to 131kb long to predict accessibility tracks. Similarly, Enformer [1] can integrate information across 200kb+, significantly improving predictions for distal regulatory elements (enhancers).

While much progress is being made, many of these algorithms were originally designed and tuned for human or mouse [10,1,23]. Models that address regulatory elements in plants are beginning to emerge, including AgroNT, which was trained on data from 48 plant species (mostly crops) [19]. It is unclear how useful these models would be for the many unrelated, non-model plant species for which little information is known about their regulatory elements and where obtaining experimental validation of regulatory elements is prohibitive. To overcome these limitations, we propose the use of co-linear chaining, a computational technique that allows for the identification of chains of small but clustered regions of similarity, such as those found within accessible chromatin regions, within and across genomes. This method combines the use of ATAC-seq peaks that highlight active regulatory elements, any available information on known regulatory motifs, and sequence similarity to predict active regulatory elements within and across genomes. To the best of our knowledge, our work is the first attempt at developing a framework for ACR prediction within genomic data using co-linear chaining.

## 3 Methods

As depicted in Figure 2, we developed a pipeline to test the utility of combining co-linear chaining with machine learning to predict chromatin accessibility. First, we obtain the sequence and ATAC-seq data of an *Arabidopsis thaliana* genome, which we use to identify ACRs. Next, we label motifs (potential CREs) from the *A. thaliana* DAPv1 database [22], which contains binding sites for hundreds of transcription factor, in the sequences. Then, we assign positive regions as the HELD-OUT set, and we assign both positive and negative regions as the EXPERIMENT set. We further split the EXPERIMENT set into TRAIN, VALIDATE, and TEST sets. We then obtain a feature vector for every region in EXPERIMENT by colinear-chaining a given region in EXPERIMENT with every region in HELD-OUT. Finally, we use these feature vectors in machine learning, masking TEST until after the model training and tuning to evaluate the accessibility predictions. Our software tool, PanACRpred, is freely available at: https://github.com/msu-alglab/pan_ACR_pred.

**Fig. 2.**
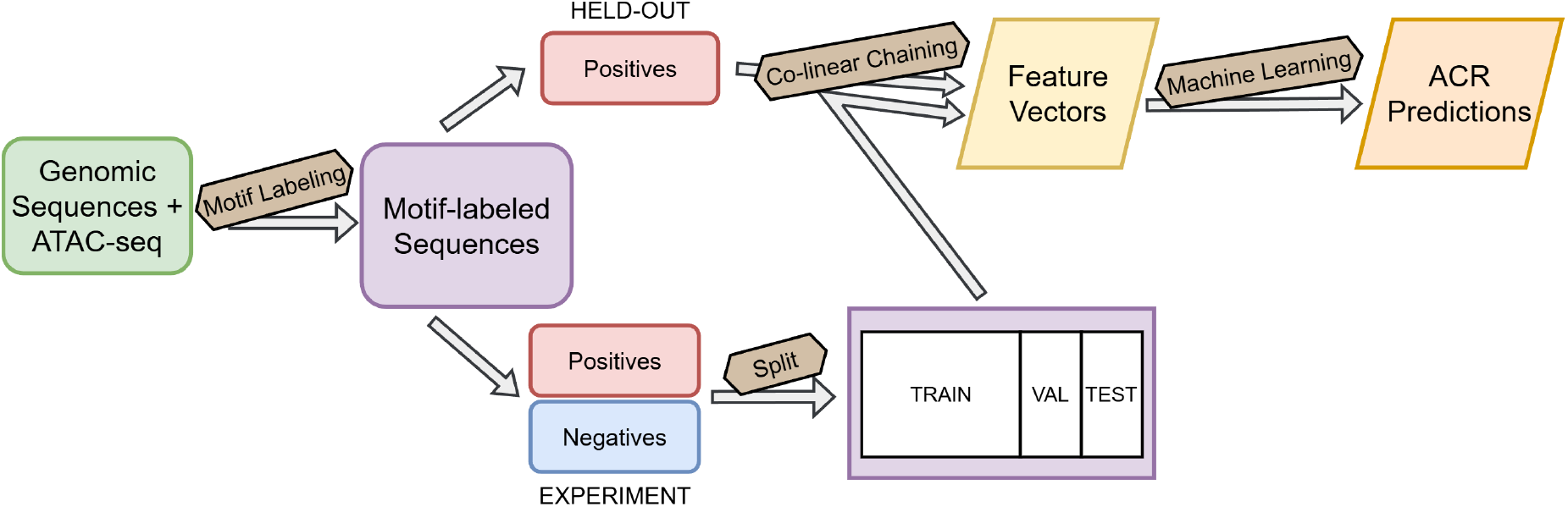
Pipeline to process genomic data and predict ACRs via co-linear chaining and machine learning.

### 3.1 Data

We obtained genome coordinates for ACRs identified in the *Arabidopsis thaliana* reference genome (Columbia-0) that differed between wildtype and MET1 mutants with reduced CG methylation across the genome [29]. We extracted the underlying sequence data from the TAIR10 assembly version using bedtool’s getfasta tool [25,12].

To obtain negative regions (inaccessible regions of the genome) for machine learning, we employ two separate generation methods: *random* and *upstream*. In the random method, we select random regions of the genome not identified as ACRs, fixing the lengths to match the length distribution of the positive ACR set used as samples for machine learning. In the upstream method, we select proximal regions not identified as ACRs upstream of known genes, once again matching the length distribution of the positive ACR set. The distance upstream from the gene is randomized, but does not exceed 2,000 base pairs.

To label motifs as potential CREs in the sequences, we used the DAPv1 database [22] of known *Arabidopsis thaliana* CREs. To avoid overcounting CREs, we clustered similar motifs in the database using a motif comparison tool, Tomtom [7], and hierarchical clustering. We then define a consensus motif for each cluster and use FIMO [6] to find these consensus motifs in the genomic sequences. Both Tomtom and FIMO are part of the MEME suite [2].

### 3.2 Co-linear Chaining

We define a string to be a sequence of characters in the alphabet *Σ* = {A, C, G, T .} A motif in a string *S* is a subsequence of the characters in *S*. For some motif *M* in *S* which begins at position *i* in *S*, let INDEX(*M*)= *i*.

Suppose we have a string *A* with a sequence of motifs *M*_*A*_ = *a*_1_ … *a*_*n*_, with INDEX(*a*_*i*_) ≤ INDEX(*a*_*i*+1_) for all *i*. Likewise, suppose we have a string *B* with a sequence of motifs *M*_*B*_ = *b*_1_ … *b*_*m*_, with INDEX(*b*_*k*_) ≤ INDEX(*b*_*k*+1_) for all *i*. Let *X* = *{*(*a*_*i*_, *b*_*k*_) | *a*_*i*_ *∈ M*_*A*_, *b*_*k*_ *∈ M*_*B*_, *a*_*i*_ = *b*_*k*_*}*. We call some element *x ∈ X* a *motif match*. Now we define a precedence relation *≺* on *X*, where (*a*_*i*_, *b*_*k*_) *≺* (*a*_*j*_, *b*_*l*_) if and only if *i < j* and *k < l*.

#### Definition 1.

*We define the* global chain length *to be the cardinality of the largest set S ⊆ X with ≺ as a total relation on S. That is, for all s, t ∈ S with sΣ*= *t, either s ≺ t or t ≺ s*.

Additionally, we can assign some weight *w*_*x*_ to the motif associated with every *x ∈ X*. For some *S ⊆ X*, let 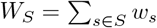

#### Definition 2.

*We define the* global chain score *to be the maximum W*_*S*_ *over all S⊆ X with≺ as a total relation on S*.

The global chain length and global chain score between two sequences with identified motif matches can be computed in 𝒪 (|*X*| log |*X* |) time with existing co-linear chaining methods methods, e.g. [17]. Fig. 3 shows the top-scoring example from the tested data set.

**Fig. 3.**
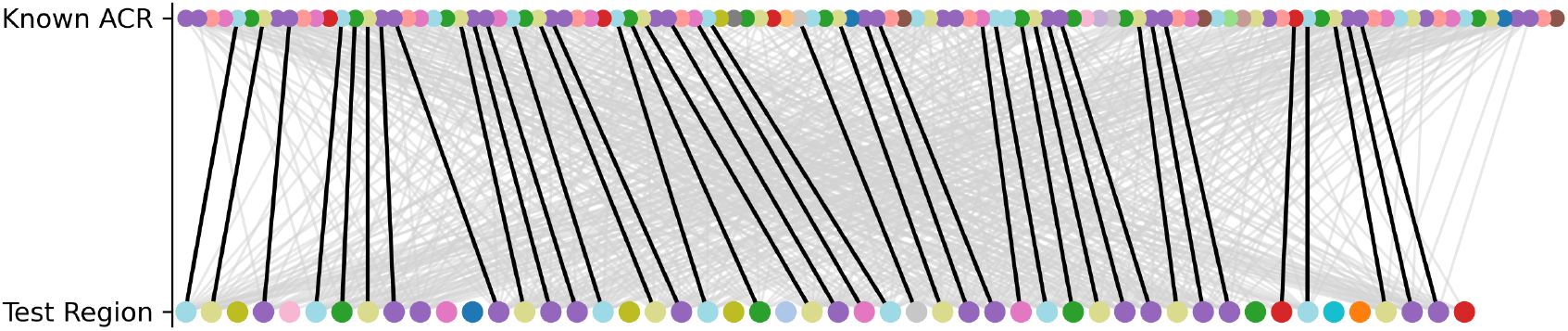
Illustration of co-linear chaining between the HELD-OUT ACR and EXPERIMENT region pair with the highest global chain score. Dots represent motifs; grey lines connect pairs of matching motifs; black lines connect pairs of matching motifs in some set which achieves the global chain score. The highest scoring pair is shown.

### 3.3 Generating Feature Vectors

Given some set *H* (the HELD-OUT set from Fig. 2) of known ACRs, with |*H*| = *n*, we can create a feature vector for some DNA region *r* of unknown accessibility. First, we arbitrarily assign an order to the ACRs in *H*. Then, we use co-linear chaining to find the global chain score *c*_*i*_ between *r* and *h*_*i*_ for every *h*_*i*_ *∈ H*. This allows us to define a feature vector *v*_*r*_ =[*c*_1_ *c*_2_ *…c*_*n*_]. This feature vector gives a quantification of the relationship between *r* and the set of known ACRs.

To calculate the global chain score, we assign a weight *w*_*m*_ to every motif *m* identified as a potential CRE. The weight is based on the frequency at which *m* appears in our set of known ACRs as opposed to the whole genome. Let prob(*m*_ACR_) be the probability that an identified motif in a known ACR is *m*. Likewise, let prob(*m*_BG_) be the probability that an identified motif in the background (in this case, the entire genome) is *m*. We calculate *w*_*m*_ as follows, setting *λ* = 0.8 based on initial experiments.

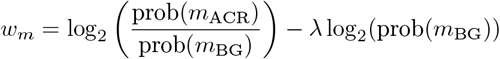

The first term in the equation scales with the frequency of a motif’s occurrence in known ACRs as opposed to the background. The second term adds an additional boost to the weight which scales with the rarity of the motif in the entire genome. The logarithmic scale ensures the weights are confined to a more narrow range. The intention behind this weighting scheme is to incorporate additional predictive power into the feature vectors by enhancing motifs which are more unique to ACRs. Thus, unknown regions which contain motifs more often found in ACRs will likely have a higher global chain score with some of the known ACRs.

### 3.4 Machine Learning Models

The main objective to predict ACRs was formulated as a supervised binary classification problem, in which instances are sequence regions, and features are the changing scores (i.e., scores between instances and sequences in the HELD-OUT set). A label of 1 indicates that the region is chromatin accessible and a label of 0 indicates that the region is not chromatin accessible. We selected the following benchmark classifiers based on preliminary testing with the data: Multi-layer Perceptron (MLP), eXtreme Gradient Boosting (XGBoost), Gradient Boosting Classifier (GBC), and Histogram Gradient Boosting Classifier (HistGBC). In addition to these classifiers, we also tested a a Naïve classifier that uses only a threshold on the chaining scores to make predictions (i.e. no machine learning) to provide a baseline comparison.

### 3.5 Experimental Setup

After labeling candidate CRE motifs in the genome, we randomly choose 12,806 ACRs as our HELD-OUT set. Next, we combine 11,758 ACRs with 11,758 negative regions to form our EXPERIMENT set. We then perform a 60% / 20% / 20% split on EXPERIMENT into TRAIN / VALIDATE / TEST sets, respectively, ensuring a 50% / 50% split between positive and negative samples in each dataset. We use co-linear chaining to obtain a feature vector for each sample, as described in Section 3.3. The results are TRAIN, VALIDATE, and TEST sets containing 14108, 4704, and 4704 samples, respectively, with each sample represented by 12,806 features.

To evaluate the performance of our models, we utilized hold-out validation, where TRAIN is used for developing the model, VALIDATE is for tuning some model parameters, and TEST is used for calculating performance metrics. We use Area Under the Receiver Operating Characteristic (AUROC) as our primary performance evaluation metric. AUROC is a key machine learning metric evaluating a model’s ability to distinguish between positive and negative classes, essentially showing its classification performance across all possible thresholds.

For MLP, we leveraged Keras/TensorFlow in addition to using Dense and Dropout layers in Python. We used 3 hidden layers with 128, 64, and 32 neurons. Dropout layers were placed between each of these hidden layers (dropout rate of 0.2 was used) and *relu* activation was used for all layers, with the exception of the output layer, which used *sigmoid* activation. This architecture was used for experiments with both the random and upstream negative sets. For GBC, HistGBC, and XGBoost, we used Scikit-learn. For GBC, we used a learning rate of 0.1 and a max depth of 3 for both the random and upstream negative set experiments. For HistGBC, we set no max depth for both the random and upstream negative set experiments. The learning rate was set to 0.1 and 0.01 for random and upstream, respectively. For XGBoost, the random negative set experiments utilized a learning rate of 0.01 and 50 estimators, while the upstream negative set experiments used a learning rate of 0.1 and 100 estimators.

In addition to these models, we implemented a custom Naïve classifier, which uses a simple, threshold-based prediction strategy. In the training step, the model identifies the average maximum chaining score over all positive samples in the train set as the threshold. Then, every test region with a maximum chaining score greater than or equal to that threshold is predicted to be an ACR.

The model hyperparameters used for evaluating the models’ performance on TEST were chosen after tuning with VALIDATE. The tuned hyperparameters are summarized in Table 1. For GBC, HistGBC, and XGBoost, we performed tuning on their learning rates, max depth, and number of estimators used via GridSearchCV (Scikit-learn). GBC saw benefit in reducing its max depth parameter while keeping its learning rate at a moderate value. This trend is not as clear for HistGBC and XGBoost, as we see more variation in the hyperparameter tuning results. For MLP, the hyperparameter tuning explored the dropout rate between layers and the number of neurons in each layer, as well as the ordering of the layers. The result was the decision to use a higher dropout rate and a layer architecture that decreases the number of neurons with the addition of layers.

**Table 1.** Hyperparameters tuned for each model.

| Classifier | Hyperparameters Tuned |
| --- | --- |
| GBC | learning rate, max depth |
| HistGBC | learning rate, max depth |
| XGBoost | learning rate, number of estimators |
| MLP | model layers / architecture, dropout rate |

## 4 Experimental Results

All classifiers (excluding the Naïve classifier) gave very similar AUROC values, differing by at most .009 when utilizing the same data (Table 2). GBC gave a slightly higher AUROC value for the experiment with the random negative set. MLP gave a slightly higher AUROC value for the experiment with the upstream negative set. The experiment with randomly generated negative regions gave higher AUROC values across all models, including the Naïve classifier (Figure 4). All classifiers perform better than the Naïve classifier by AUROC value and other performance metrics, as summarized in Table 2.

**Table 2.**
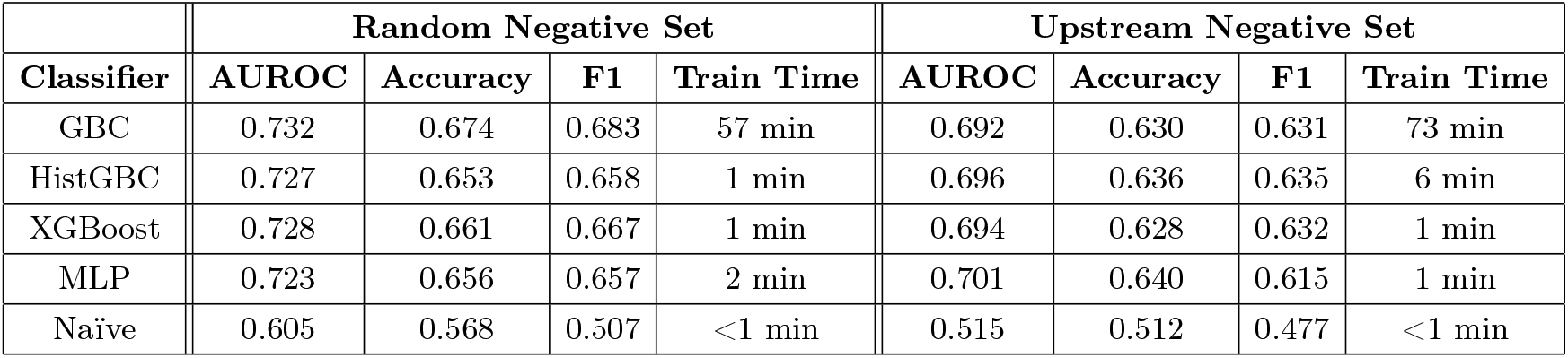
Summary of various models’ performance across experiments with both types of negative regions.

| Classifier | Random Negative Set |  |  |  | Upstream Negative Set |  |  |  |
| --- | --- | --- | --- | --- | --- | --- | --- | --- |
|  | AUROC | Accuracy | F1 | Train Time | AUROC | Accuracy | F1 | Train Time |
| GBC | 0.732 | 0.674 | 0.683 | 57 min | 0.692 | 0.630 | 0.631 | 73 min |
| HistGBC | 0.727 | 0.653 | 0.658 | 1 min | 0.696 | 0.636 | 0.635 | 6 min |
| XGBoost | 0.728 | 0.661 | 0.667 | 1 min | 0.694 | 0.628 | 0.632 | 1 min |
| MLP | 0.723 | 0.656 | 0.657 | 2 min | 0.701 | 0.640 | 0.615 | 1 min |
| Naïve | 0.605 | 0.568 | 0.507 | <1 min | 0.515 | 0.512 | 0.477 | <1 min |

**Fig. 4.**
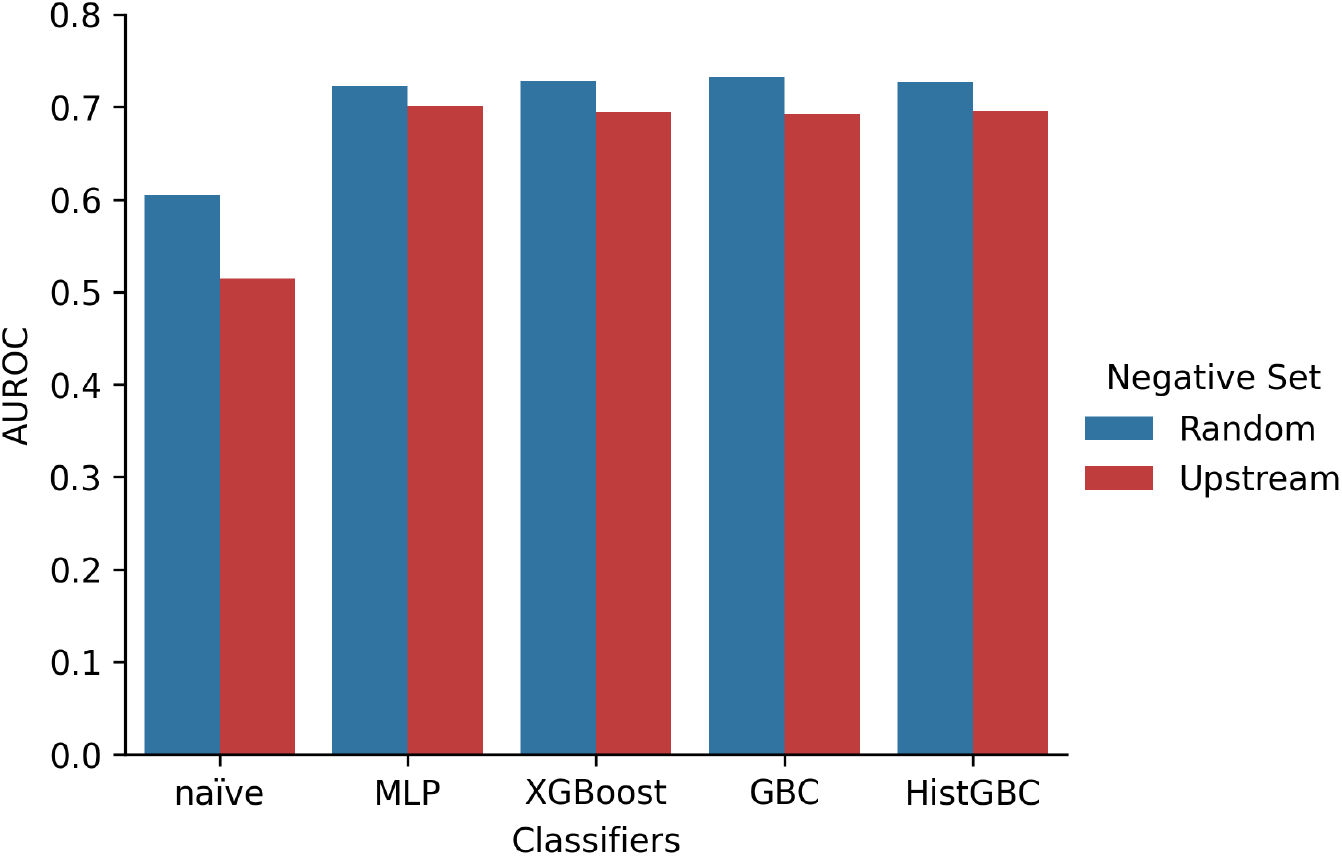
Bar graph showing the classifier performances based on AUROC values. AUROC (Area Under the Receiver Operating Characteristic Curve) evaluates a model’s ability to distinguish between positive and negative classes. It’s a single number (0 to 1) summarizing the ROC curve, where 1 is perfect, and 0.5 is random guessing. Negative set explains the methodology for generating the negative samples for the data e.g. Random negative set describes the generation of negative samples via random selection.

The runtimes to train GBC, HistGBC, and XGBoost were under 7 minutes for all experiments and were under 2 minutes in most cases. GBC train times, on the other hand, were significantly longer, at 57 minutes for the random negative set experiment and 73 minutes for the upstream negative set experiment (Table 2). All timing data was collected on Google Colab Pro with an AMD EPYC 7B12 processer (2.25 GHz), equipped with 4 cores and 50 GB of available RAM.

## 5 Discussion

Here, we have shown that the conservation of clusters of short regulatory motifs in conjunction with ML models can be used to accurately predict ACRs, identifying genomic regions that are likely to be important in controlling gene expression and the resulting phenotypes. Our ultimate goal is to predict ACRs across multiple genomes within a pangenome, identify both known and novel motifs within the ACRs, and use these as features to predict phenotypes and better understand which genomic variants are driving important phenotypic traits.

In the future, this project will also include the implementation of Positive Unlabeled (PU) learning. In our current work, the gold-standard data is composed of positive examples (i.e., known ACR regions), and we had to develop methods to generate our negative examples. Because our positive data is incomplete, the manual generation of negatively labeled samples may include examples that are positive in other tissues or conditions, giving rise to concerns about whether all our negative data are truly negative. However, with the use of PU learning in future work, we can explore its ability to circumvent this underlying concern with our data by considering the performance results from a dataset consisting of only positively labeled or unlabeled samples.

*Acknowledgments* Support for this project came from NSF awards 2414134 and 2243010.

